# Analysis of Seasonal and Long-Term Population Dynamics for Modeling Populations at Low Density: Experience with Light Traps

**DOI:** 10.64898/2026.03.23.713576

**Authors:** Vladislav Soukhovolsky, Vladimir Dubatolov, Anton Kovalev, Olga Tarasova, Vyacheslav Martemyanov

## Abstract

Methods for estimating and modeling the long-term and short-term adult flight dynamics of the conifer silk moth *Dendrolimus superans* (Lepidoptera: Lasiocampidae) are examined. The analysis uses light trap adult catch data collected over 21 years, from 2005 to 2025. Three models of adult flight are considered: a flight-initiation model driven by weather factors, an autoregressive model of long-term catch dynamics, and a binary model of seasonal catch. For the flight-initiation model, we propose estimating the accumulated temperature sum ST from the date when the first derivative of the remote sensing vegetation index NDVI becomes positive until the date of the first adult capture of the season. ST is shown to be sufficiently stable across all years of observation, with flight each year beginning after this temperature sum is reached. The second model demonstrates that the long-term light trap catch time series is well described by a second-order autoregressive model AR(2), in which the catch of the current year depends on catches from the two preceding years. This long-term series is compared with a previously studied larval population density series of the Siberian silk moth; both are shown to be AR(2) series with similar coefficient values, which suggesting that adult catch data may serve as a proxy for absolute larval population density. In the third model, we describe the transition from absolute-scale seasonal catch dynamics (number of adults per day) to a binary scale (0, 1), where 0 denotes days on which no adults were attracted to the trap, and 1 denotes days on which at least one individual was captured. The seasonal absolute catch series is thereby transformed into a binary series of zeros and ones, and relationships between adjacent values in such a binary series are examined. A linear relationship between the absolute and binary seasonal dynamics series is demonstrated, making it possible to estimate absolute catches from binary catch values and to analyze seasonal flight in sparse pest populations. This potentially opens new avenues for understanding how outbreak populations function at chronically low density.

**Author summary:** Forest pests can cause catastrophic damage, yet predicting their outbreaks remains challenging. During periods of low population density, standard monitoring methods become labor-intensive and uninformative, while the transition to an outbreak often occurs unexpectedly.

Using a 21-year dataset of adult Siberian silk moth (Dendrolimus superans) captures from light traps, we developed an approach combining three complementary models. First, we showed that moth flight begins upon reaching a specific temperature sum, with the starting point determined by NDVI vegetation index dynamics rather than a calendar date—making the forecast more ecologically relevant. Second, long-term adult population dynamics follow a second-order autoregressive model AR(2), matching the dynamics previously observed for larval populations.

This establishes light trap data as a reliable proxy for absolute population density when ground surveys are impractical. Third, we introduced a method to analyze seasonal flight using binary data (presence/absence of moths per day), which we showed is linearly related to absolute abundance. This enables studying population dynamics during periods of extremely low density, when traditional methods fail.

Our approach opens new possibilities for early warning systems to detect when a population risks transitioning from a latent state to an outbreak phase.

## Introduction

Forecasting population outbreaks of eruptive pest species is an important goal of both fundamental and applied ecology (Royama, 1984; Fischbein & Corley, 2022). One productive approach to ecological insect monitoring is pheromone trapping, which, with relatively modest observer effort, provides a quantitative estimate of the current population density of the target species (Witzgall et al., 2010). An alternative is light trapping (Shimoda & Honda, 2013); however, this approach requires additional expertise in entomological taxonomy, since light traps for nocturnal insects (e.g., moths) have lower species specificity. This limitation is increasingly being offset by the rapid development of artificial intelligence and deep learning methods for species identification (Li et al., 2021; Teixeira et al., 2023). One of the pressing challenges in using any type of observation, including light trapping, is the low information content of time series when analyzing population density during the post-outbreak decline phase, or in stably sparse populations (Martin et al., 2005).

In our study, we assembled a 21-year time series of population density observations for *Dendrolimus superans* to characterize its population in the Russian Far East. *Dendrolimus superans* (Lepidoptera: Lasiocampidae) is a serious pest of coniferous forests in the Russian Far East and northern China (Kondakov, 1974; Isaev et al., 2001; Kharuk & Antamoshkina, 2017). For many years it was believed that the principal damage in Siberian forests was caused by the subspecies *Dendrolimus superans sibiricus* Tschetv. However, recent genetic analysis has shown that Far Eastern populations of *D. superans* form a distinct cluster, and *D. sibiricus* might now recognized as a separate species (Kononov et al., 2016), although phenotypic differences between these groups are not strongly pronounced and some authors still regard them as subspecies (Zolotukhin, 2015). Both *Dendrolimus superans sibiricus* and *Dendrolimus superans superans* may occur in the Far Eastern region. Our goal was not to assess *Dendrolimus* speciation, which, based on genetic evidence, is most probably ongoing (Kononov et al., 2016; Stewart et al., 2023; Shipova et al., 2026); we therefore do not focus on this question in the present study, noting only that in all likelihood we were working with a population of *D. superans* inhabiting the area closer to the Pacific coast.

Interest in studying members of the genus *Dendrolimus* is driven by two concerns: the risk of irreversible forest damage within the species’ range (Baranchikov & Kondakov, 1997; Kirichenko et al., 2024) and the risk of invasion into new territories (Kirichenko et al., 2008; Flø et al., 2020; Stewart et al., 2023). Given the potential risks associated with introduction of this pest into European countries, it has been included in the A2 quarantine list of EPPO (European and Mediterranean Plant Protection Organization), and stringent regulations are enforced to prevent its inadvertent introduction [EPPO]. Despite the economic importance of this pest to forestry on a global scale, information on the long-term population dynamics of this species is extremely limited. Only two sufficiently long datasets are known, presented in Kondakov (1974) and Yurchenko & Turova (2007). The scarcity of available data on such an important pest stems from two difficulties in population censusing. First, monitoring typically begins when the pest enters the outbreak phase and, by damaging trees, effectively marks the zone where reliable counts can be conducted. Second, once the outbreak ends, the pest disappears from the focus area, and the appropriate location for further surveys becomes unclear. In the above-cited studies (Kondakov, 1974; Yurchenko & Turova, 2007), surveys were conducted across a sufficiently large territory, and the published data can be regarded as integrated population characteristics. Long-term monitoring of pest populations in a stably sparse state is problematic, since it is unclear where to conduct surveys and what the practical significance of such monitoring would be in a territory where outbreaks may never occur. However, studying pest population dynamics at low density may yield important information about the regulatory mechanisms of the species.

Since no stand damage occurs at low population density, damage data cannot be used to select survey sites, and locating larvae or pupae across vast taiga forest areas is impractical. Only one method can be proposed for studying the long-term dynamics of the Siberian silk moth at low density: long-term monitoring of adult catches using pheromone or light traps.

If a light trap is regarded as a tool for estimating pest population density, there are two approaches to measuring trap catch: cumulative and differential. In the cumulative approach, the trap is placed at a site at time t_0_, approximately before the onset of adult flight, and the total catch is counted at approximately the time of flight cessation t_f_. In the differential approach, the trap is placed at time t_0_ and catch is recorded at intervals of every m days (at the limit, m = 1, i.e., daily catch is recorded). The differential method is considerably more labor-intensive; however, whether the additional information on population state justifies this increased effort remains an open question. In the present study, data on adult catches by light trap at a local site with coordinates 48.2985°N, 134.8222°E in the forests of the Russian Far East about 20 km west from Khabarovsk from 2005 to 2025 were used to analyze seasonal adult flight dynamics and long-term population dynamics of *Dendrolimus superans*.

### Study Organisms and Study Site

Figure 1 shows a satellite image of the study area, and Table 1 presents the coordinates of the light trap and study plots as potential sources of pest adults.

**Table 1.** Location of the light trap and conifer (host plant) stands P01–P03 near the village of Bychiha (Russian Far East) as sources of Siberian silk moth adults.

| Study plot | Coordinates |  | Distance from trap to plot (m) |
| --- | --- | --- | --- |
|  | Latitude (°N) | Longitude (°E) |  |
| Light trap | 48.2985 | 134.8222 | 0 |
| P01 | 48.2838 | 134.8127 | 1790 |
| P02 | 48.2766 | 134.8169 | 2480 |
| P03 | 48.2706 | 134.8166 | 3145 |

**Figure 1.**
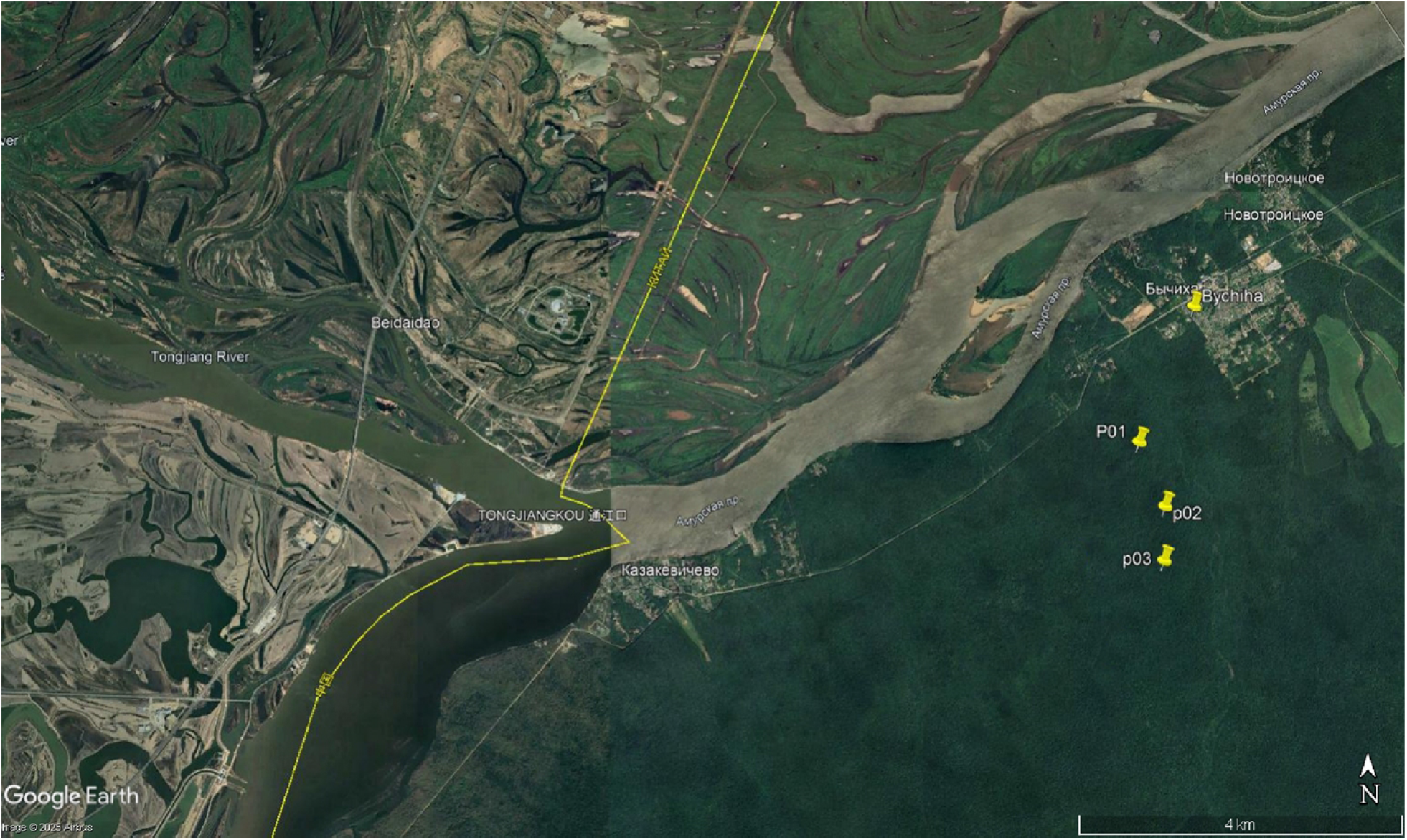
Satellite image of the study area.

## Materials and Methods

### 1. Ground-Based Adult Flight Monitoring

Ground-based moth flight monitoring of *D. superans* was conducted at the outskirts of the village of Bychiha (48.2985°N; 134.8222°E) at the edge of a mixed coniferous–broadleaved forest. A mercury-quartz fluorescent DRV lamp (160 W, luminous flux 2300 lm).

### 2. Assessment of Stand Condition and Potential Tree Damage

To assess potential foliage removal by insects, the seasonal dynamics of the photosynthetic vegetation index NDVI were examined. It is known that when trees are damaged by folivorous insects, both the seasonal NDVI sum and the seasonal NDVI maximum decrease (Kovalev et al., 2024). Accordingly, changes in the seasonal NDVI curve can serve as an indicator of substantial damage levels. Seasonal NDVI data were obtained from the MODIS AQUA/TERRA satellite system operating under NASA’s Earth Observing System (EOS) program (Barnes et al., 1998). These data were used to calculate derived indices: the seasonal NDVI sum and the seasonal NDVI maximum.

### 3. Model for the Onset of Adult Flight

We propose estimating the conditions at which adult flight to the light trap begins following the approach used to assess phenological phase duration in trees.

Leaf budburst in forest stands is forecast as follows. A reference start date t_0_ is selected corresponding to the day when daily temperature crosses a critical value T_0_ (typically +5 °C). The date t_1_ of leaf budburst onset is then determined from phenological observations. Knowing these two dates, the accumulated temperature sum ST for all days between t_0_ and t_1_ is calculated from meteorological data. In theory, ST should be constant across all years. Having calculated ST from a set of seasons, the accumulated temperature sum from t_0_ to an arbitrary date t_j_ can be computed for any given year, allowing the onset of the phenological phase to be forecast (Phenology, 2013; Wolkovich et al., 2014; Delpierre et al., 2016).

The same approach can be used to estimate the onset date of adult flight. Three parameters are required: the date t_0_ at which temperature crosses the critical value T_c_, the date t_1_ of the first adult capture, and daily air temperature data for the intervening period. Phenological studies typically rely on meteorological station data; however, given the very sparse station network in the Asian part of Russia, the nearest station to the observation site may be more than 100 km away, making such data unrepresentative. Therefore, daily satellite land surface temperature (LST) data were used to estimate local air temperature in the study area. The satellite LST pixel size is 1 × 1 km, which is entirely adequate for reliable estimation of local weather conditions.

### 4. Autoregressive Model of Long-Term Catch Dynamics

The authors have previously shown that an autoregressive model AR(k) can be used to describe long-term insect population dynamics, where the current population density depends on densities from the preceding k seasons (Soukhovolsky et al., 2015). In general form, the AR(k) model for long-term catch dynamics is expressed as:

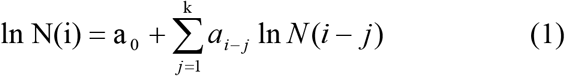

If the order of autoregression and the values of the time series {N(i)} in equation (1) are known, equation (1) can be treated as a linear regression equation with unknown parameters a_0_, …, a□, whose values can be estimated using standard linear regression methods (Wei, 2006).

### 5. Binary Model of Seasonal Catch Dynamics

Flight events during the season can be divided into two types. **Regular** flight is characterized by a nonlinear unimodal temporal dependence of flight intensity, with daily catch substantially greater than zero. **Sparse** flight is characterized by the absence of daily adult captures and low absolute catch.

To describe flight dynamics as a set of rare events, we convert the absolute daily catch scale (number of adults per day) to a binary scale, where 0 denotes days on which no adults were attracted to the trap, and 1 denotes days on which captures occurred. Flight dynamics are thereby represented as a binary series of zeros and ones.

For such a series, the presence of a temporal trend – manifested as a change in the probability of observing state 0 over time – can be assessed. If for different time segments p(0) = const, the binary time series can be regarded as stationary with probability p(0) of observing state 0 and probability p(1) of observing state 1. Clearly, p(0) + p(1) = 1.

A key question is whether the binary series is independent, and if not, whether the probability p(0, j) at time j depends on the state of the series at times (j-1), (j-2), … (j-k).

In the simplest case, where the state of the series at time j depends on its state at time (j−1), we are dealing with a Markov chain (Wai-Ki Ching et al., 2013).

Testing the Markov properties of the series is straightforward: the numbers of event pairs (0,0), (0,1), (1,0), and (1,1) are calculated for the series under study, and these are compared with the expected counts [p(k)×p(r)×N] using the χ^2^-test, where p(k) and p(r) are the proportions of events with values k = 0,…1 and r = 0,…1, and N is the length of the time series. If the χ^2^-test value is less than the critical value for the given degrees of freedom, the observed series can be said to lack Markov properties, and the current value of the series is independent of the preceding value.

## Results

### 1. Flight-Initiation Model

Since there is no theoretical basis for selecting the reference date t_0_ from which accumulated temperature is calculated, t_0_ was determined from seasonal NDVI dynamics. Two time markers were proposed: the first reference point t_0_ – the date when NDVI begins to rise in spring and the difference between successive NDVI values becomes positive; the second point t_1_ – the day when the first adult is found in the light trap. The accumulated temperature sums SLST between points t_0_ and t_1_ at the trap location should be constant across years for a given species following the same logic as heat sums used for phenological phases.

For the proposed method, the critical points t_0_ and t_1_ were determined (typical seasonal NDVI curves and first-difference NDVI series are presented in Figure 2).

**Figure 2.**
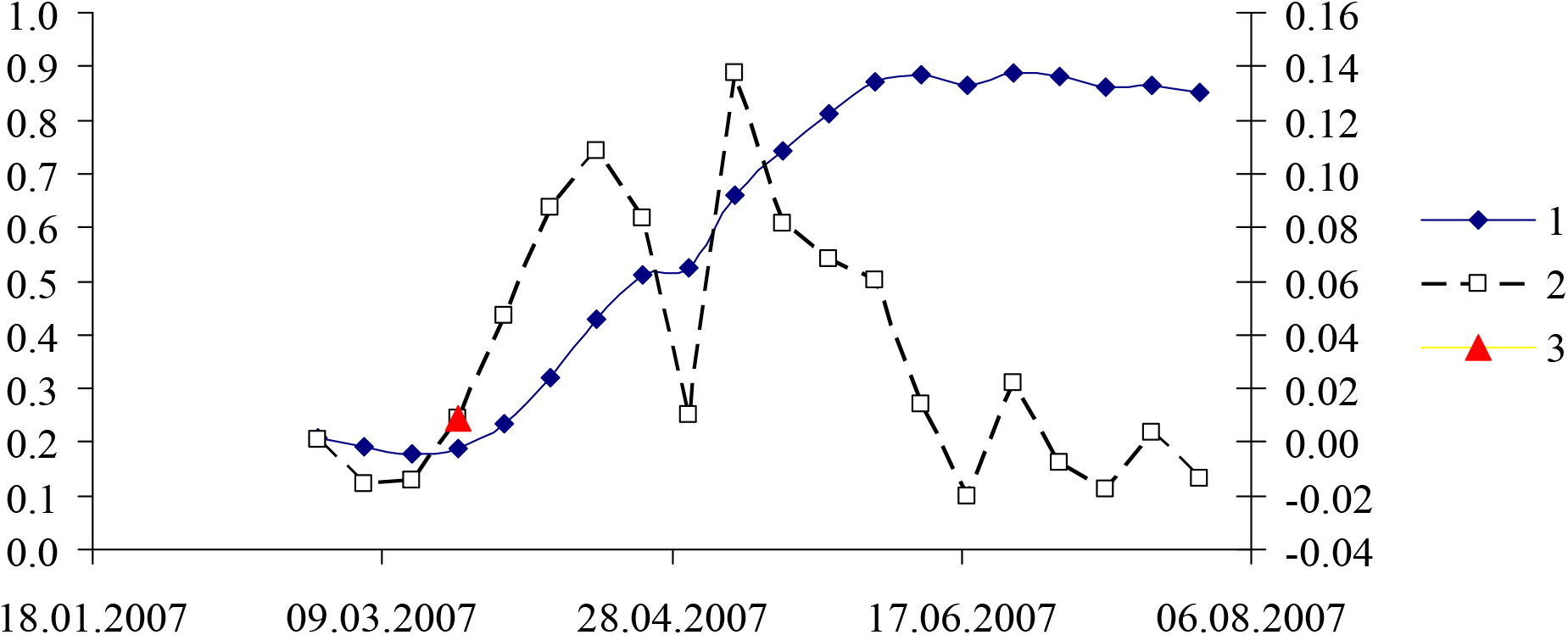
Seasonal dynamics of the NDVI series (1), the first-difference NDVI series (2), and the critical point (3).

For each year, the logarithm of the daily LST sum from date t_0_ to date t_1_ was calculated (Table S1 in the Supplement). Calculations showed that the values of SLST(i) for all i (i = 1, …, 21) years are very similar: the long-term mean SLST equals 7.38, with a standard deviation s = 0.23 and standard error s_x_ = 0.054. The variability of SLST(i) among individual years is therefore very small, and flight begins each year after the accumulation of an approximately constant temperature sum.

### 2. Long-Term Catch Dynamics Model

Assuming the long-term dynamics series is described by an autoregressive model of equation (1), the order k of the AR(k) model must first be determined The standard approach is to calculate the partial autocorrelation function (PACF) (Box & Jenkins, 1970). The autoregressive order is defined as the value of k at which the PACF falls within its standard error bounds. The PACF for the long-term adult catch series of the Siberian silk moth is shown in Figure 3.

**Figure 3.**
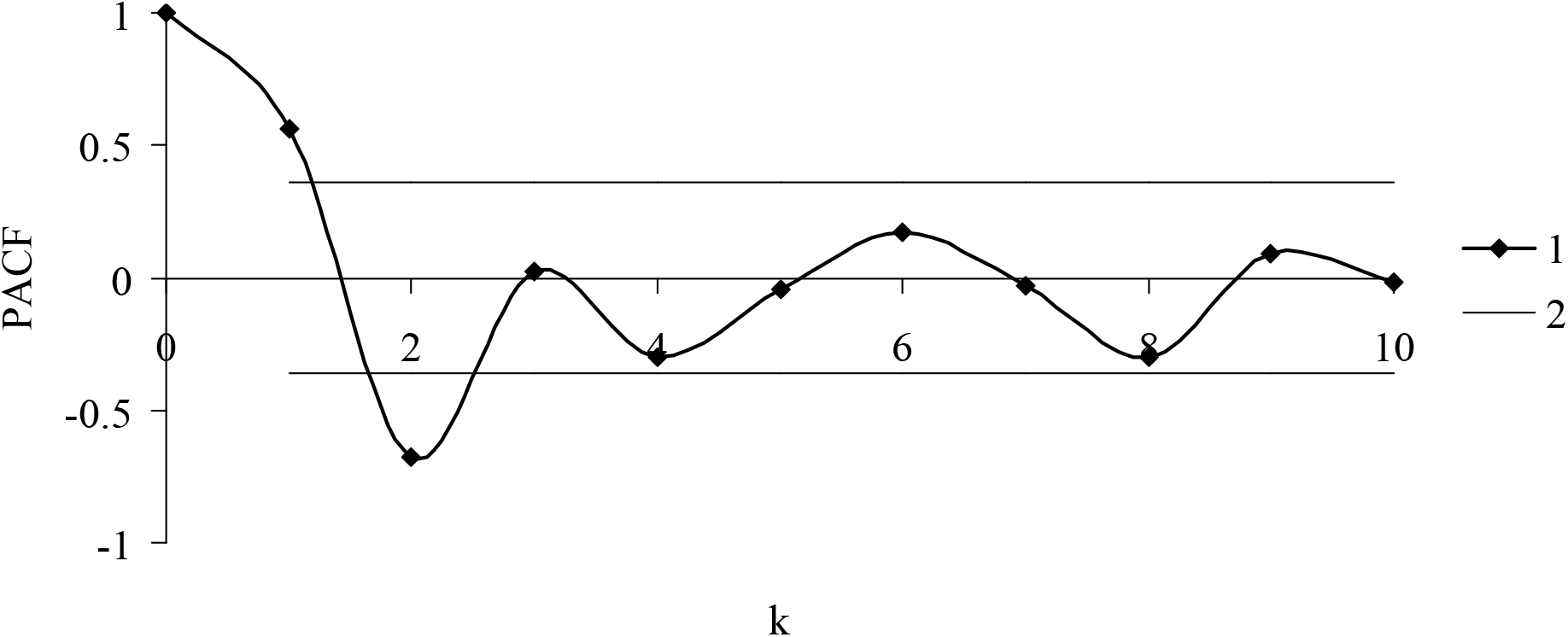
Partial autocorrelation function (PACF) of the long-term catch dynamics series. 1 – PACF; 2 – std. error of PACF.

As shown in Figure 3, the autoregressive order k equals 2 for the catch series. The AR(2) model coefficients are given in Table 2.

**Table 2.** AR(2) model for the catch series.

| Parameter | Coeff. | Std.<br>Err. | t-test | p-value |
| --- | --- | --- | --- | --- |
| a0 | 14.415 | 4.558 | 3.162 | 0.008183 |
| N(i-2) | -0.732 | 0.196 | -3.732 | 0.002863 |
| N(i-1) | 0.997 | 0.199 | 5.018 | 0.000300 |
| Adj R<sup>2</sup> | 0.630 |  |  |  |
| F-test | 13.14 |  |  |  |

Figure 4 shows the time series of seasonal catch N(i) over the study period. To reduce random errors, high-frequency filtering was applied using the Hann filter:

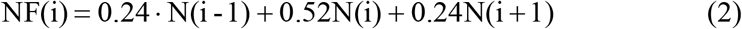

**Figure 4.**
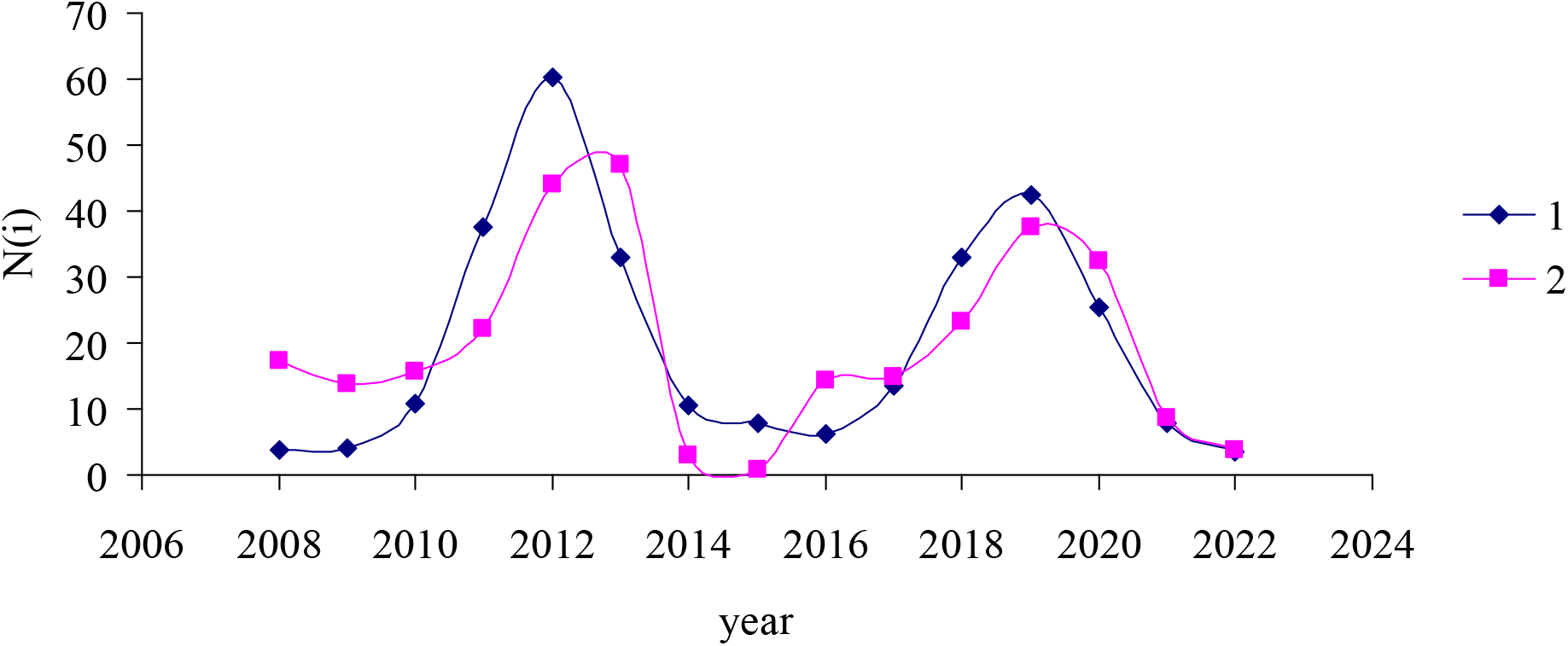
Long-term catch dynamics of the Siberian silk moth after high-frequency filtering with the Hann filter. 1 – observed data; 2 – AR model.

As seen in Table 2, the current light trap catch is influenced by the catches of seasons (i-1) and (i-2). The coefficient a_1_ is positive and a_2_ is negative. These coefficients describe how current catch is regulated by catches from the two preceding years. The same signs of the feedback coefficients are characteristic of population dynamics series for the Siberian silk moth and, more generally, of a large number of forest insect species exhibiting outbreak dynamics (Soukhovolsky et al., 2025). For comparison, the larval density series for the Siberian silk moth in the Russian Far East is described by the model equations in Table 3.

**Table 3.**
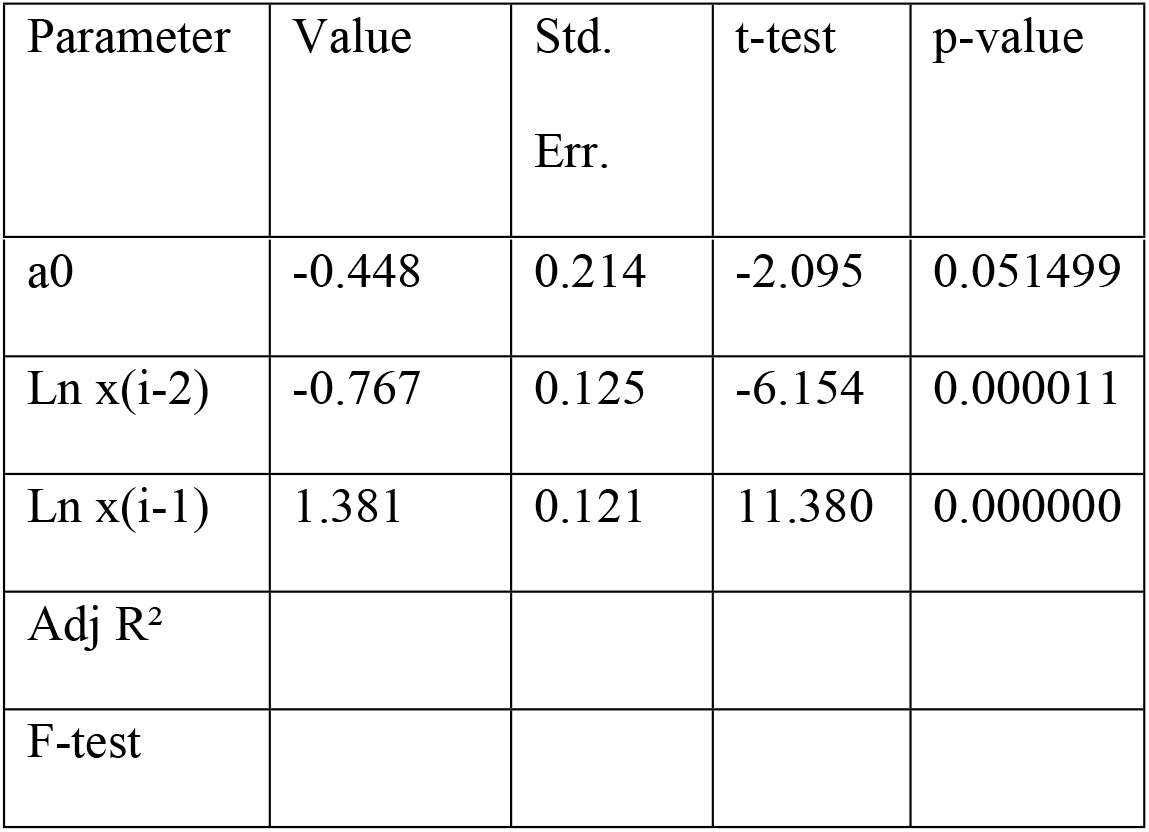
AR(2) model coefficients for the larval density series x(i) of the Siberian silk moth in the Russian Far East.

Figure 5 shows field observation data from Yurchenko & Turova (2007) and the modeled AR(2) population density series for the Siberian silk moth in Khabarovsk suburbs.

**Figure 5.**
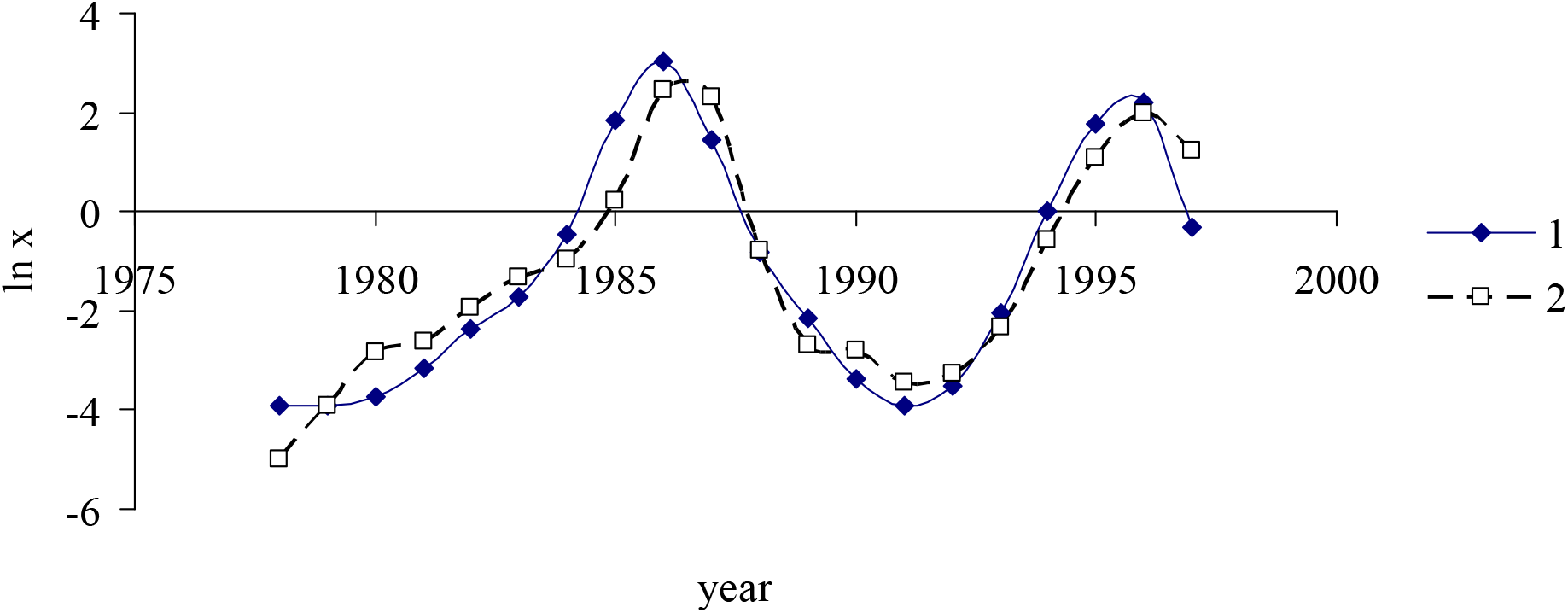
Population dynamics of the Siberian silk moth in the Russian Far East. 1 – observed data (Yurchenko & Turova, 2007); 2 – AR(2) model (after Soukhovolsky et al., 2025).

Comparing Figures 4 and 5, it can be concluded that although the series differ in developmental stage assessed (larvae vs. adults) and the analyzed period (adults: 2005–2025; larvae: 1978– 1997), the model coefficients are very similar. This indicates that the oscillation periods of the larval and adult count time series are comparable.

Does weather during the flight period and stand condition influence light trap catch? As a weather indicator, we use the mean LST value during the flight period, and as an indicator of stand condition near the trap – the response function characterizing the NDVI response to changes in LST. The correlation coefficient between the mean seasonal LST and light trap catch equals 0.03, indicating no significant relationship between weather during the flight season and trap catch.

### 3. Stand Condition and Damage Risk Assessment

When estimating larval pest population density, the risk of stand damage can be assessed if the individual consumption rate is known. However, light trap data do not allow direct estimation of absolute larval population density. The coefficient H linking seasonal light trap catch to larval population density is likewise unknown. To assess the risk of stand damage, it is therefore necessary to evaluate the relationship between total seasonal catch and the level of pest impact on the stand, as characterized by the seasonal NDVI sum and NDVI maximum. In the absence of damage, the seasonal NDVI sum should be close to 20 and the NDVI maximum close to 1. Table S2 presents long-term dynamics of the NDVI sum and maximum at study plot P01 from weeks 13 to 41. As shown in Table S2, the calculated NDVI values did not change significantly throughout the entire study period, indicating the absence of severe foliage damage by the pest. This is corroborated by our long-term field observations. Similar results were obtained for study plots P02–P05. It can therefore be concluded that no sharp increase in Siberian silk moth abundance occurred at any point during the study period, and the population remained in a stably sparse state throughout the study period.

### 4. Binary Catch Model

Table 4 presents light trap catch data for 2018.

**Table 4.** Absolute and binary light trap catches in 2018.

| Date | Absolute catch | Binary catch | Date | Absolute catch | Binary catch |
| --- | --- | --- | --- | --- | --- |
| 01.07.2018 | 4 | 1 | 16.07.2018 | 2 | 1 |
| 02.07.2018 | 1 | 1 | 17.07.2018 | 3 | 1 |
| 03.07.2018 | 0 | 0 | 18.07.2018 | 0 | 0 |
| 04.07.2018 | 1 | 1 | 19.07.2018 | 3 | 1 |
| 05.07.2018 | 0 | 0 | 20.07.2018 | 0 | 0 |
| 06.07.2018 | 0 | 0 | 21.07.2018 | 0 | 0 |
| 07.07.2018 | 0 | 0 | 22.07.2018 | 4 | 1 |
| 08.07.2018 | 1 | 1 | 23.07.2018 | 0 | 0 |
| 09.07.2018 | 0 | 0 | 24.07.2018 | 0 | 0 |
| 10.07.2018 | 2 | 1 | 25.07.2018 | 2 | 1 |
| 11.07.2018 | 1 | 1 | 26.07.2018 | 1 | 1 |
| 12.07.2018 | 0 | 0 | 27.07.2018 | 0 | 0 |
| 13.07.2018 | 0 | 0 | 28.07.2018 | 0 | 0 |
| 14.07.2018 | 0 | 0 | 29.07.2018 | 3 | 1 |
| 15.07.2018 | 3 | 1 | 30.07.2018 | 0 | 0 |
|  |  |  | 31.07.2018 | 2 | 1 |

As follows from Table 4, 33 adults were captured over 31 flight days, and the probabilities p(0) = 0.516 and p(1) = 0.484 can be readily calculated.

Table 5 presents the numbers of event pairs (0,0), (0,1), (1,0), and (1,1) for the observed binary series and for the independent series.

**Table 5.** Numbers of event pairs (0,0), (0,1), (1,0), and (1,1) for the observed binary series and for the independent series.

| Binary variable<br>value | Series type | Binary variable value |  |
| --- | --- | --- | --- |
|  |  | 0 | 1 |
| 0 | Observed | 7 | 9 |
|  | Independent | 7.99 | 7.49 |
| 1 | Observed | 9 | 5 |
|  | Independent | 7.49 | 7.02 |

From the data in Table 5, the χ^2^-test value can be calculated as:

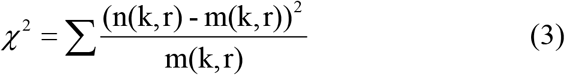

where n(k,r) is the observed number of pairs with k and r equal to 0 or 1, and m(k,r) is the expected count under the independence model.

For the data in Table 5, χ^2^ = 1.31, which is substantially less than the critical value χ^2^(cr) = 3.84 at p = 0.05. Adult flight to the light trap in 2018 can therefore be regarded as a random process with approximately equal probabilities of capture and non-capture on any given day.

Analogous data for 2019 are presented in Tables S3 and S4, and for 2020 in Tables S5 and S6. In 2019, 55 individuals were captured, p(1) = 0.55, p(0) = 0.45, and χ^2^ = 0.48, indicating that connections between adjacent values in the binary series are insignificant (p = 0.49 И0.42). In 2020, 21 individuals were captured, p(1) = p(0) = 0.50, and χ^2^ = 0.65, which indicates the independence of adjacent values in the contingency table S6.

As flight transitions from sparse random to regular, p(1) → 1. In the case of very sparse random flight, p(0) → 1 (i.e., no individuals were captured). Flight type can therefore be classified by the parameters p(1) and p(0). For the 2018–2020 data, p(1) ≈ p(0) ≈ 0.50, indicating that the flight type was approximately the same across all three years.

The ecological causes of flight as a rare event primarily reflect low insect density in the study area, although poor propagation of the light signal cannot be excluded, since light travels in straight lines and landscape elements may obstruct its interaction with moths (scattering, absorption, etc.). In such cases, it is advisable to install multiple traps and to assess the variability of p(1) across them. If the maximum p(1) values for a group of traps are low, this most likely indicates low population density in the area. A trap network may also help delineate the boundaries of an outbreak zone. Another cause of intermittent flight may be weather during the flight period; however, at high population density, catches on days favorable for flight will be correspondingly high.

Analogously to the long-term absolute catch dynamics series (Figure 4), an AR(2) model of the p_1_ series can be constructed (Figure 6).

**Figure 6.**
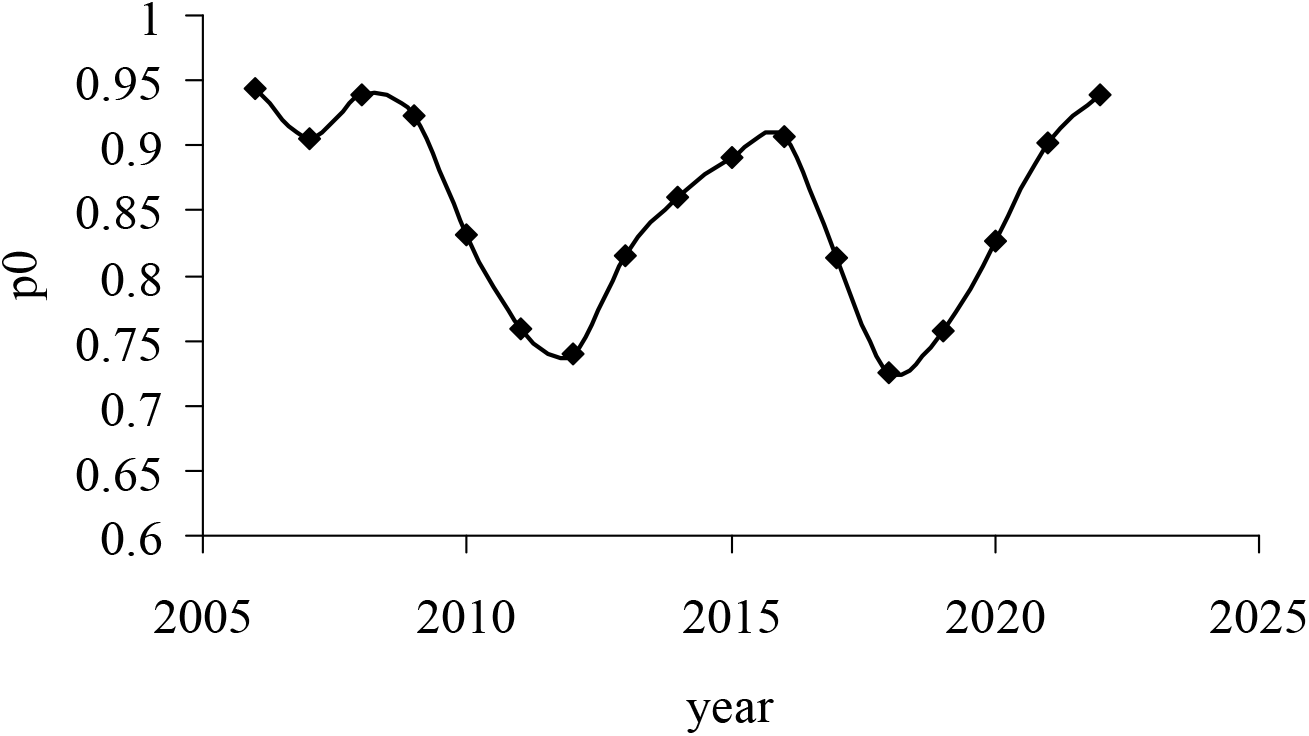
Temporal dynamics of the p_1_ series.

The order of the PACF of the p_1_ series equals 2; the estimated AR(2) model coefficients for the binary catch time series are presented in Table 6.

**Table 6.** AR(2) model coefficients for the variable p_1_.

| Parameter | Value | Std. Err. | t-test | p-value |
| --- | --- | --- | --- | --- |
| h0 | 0.496049 | 0.142827 | 3.47308 | 0.004605 |
| p0(i-2) | -0.733549 | 0.193838 | -3.78434 | 0.002603 |
| p0(i-1) | 1.147854 | 0.202945 | 5.65598 | 0.000106 |
| Adj R<sup>2</sup> | 0.680000 |  |  |  |
| F-test | 16.020000 |  |  |  |

The absolute and binary catch series are in phase, and binary catch values can be used in place of absolute catch measurements (Figure 7).

**Figure 7.**
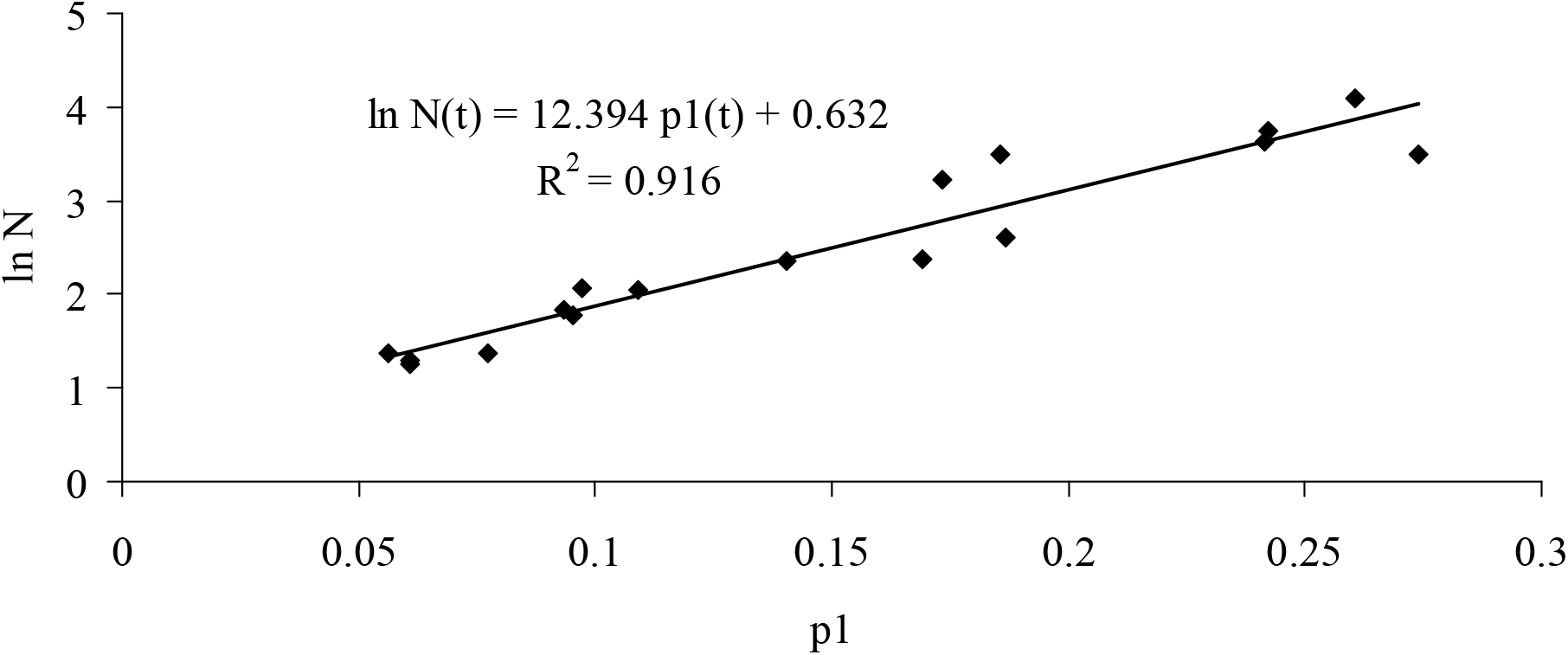
Relationship between p_1_(t) and ln N(t).

As shown in Figure 7, knowledge of p_1_ allows estimation of the absolute catch N.

## Discussion

When studying the long-term population dynamics of pest species in which periodic outbreaks lead to sharp increases in population density, the long-term dynamics curve is often cyclic. However, in most cases it remains unclear how population dynamics behave during the chronic low-density (inter-outbreak) phase, when damage foci are absent. Two types of dynamics are possible: random fluctuations around a low mean value, or cyclic oscillations similar to those during outbreaks but with a smaller amplitude. Our ground-based and remote sensing observations of the *D. superans* in the Russian Far East revealed a prolonged depression of this species in the region, consistent with data from other researchers obtained during the same period (Kirichenko et al., 2024). This was confirmed both by light attraction monitoring and by satellite data analyzed using the remote sensing approaches developed in our earlier work (Kovalev et al., 2024). The interval between population peaks was approximately 7 years, consistent with the periodicity of populations reaching peak densities (Soukhovolsky et al., 2024). Conditions in the Far East apparently do not favor the full expression of the Siberian silk moth’s outbreak potential. One possible reason is the relatively high tree species diversity in Far Eastern forests compared with the continental Siberian taiga. Being a pronounced oligophage (Rozhkov, 1963), D. superans is significantly limited in its choice of a diverse resource for the maximum realization of biotic potential and increase in population density.

Comparison of the model parameters in Table 2 with those of the model previously developed for long-term Siberian silk moth dynamics in the forests of Khabarovsk Krai (Soukhovolsky et al., 2025) shows that both outbreak populations and populations at low density are governed by the same system of positive and negative feedback. An outbreak may therefore not involve the emergence of a new regulatory mechanism but rather reflect, for example, a change in the state or availability of food resources (Soukhovolsky et al., 2023). It can accordingly be hypothesized that at low population density, regulatory processes operate through the same mechanism that governs fluctuations during outbreaks.

A key finding of the modeling component is that binary models enable analysis of time series with numerous zero-catch observations to evaluate seasonal flight dynamics. After accounting for accumulated temperature sums, flight onset timing was shown to be directly governed by ambient temperature conditions. The Siberian silk moth exhibits ontogenetic variability. Usually, eggs are laid on host tree needles, and first-instar larvae feed for about 3–4 weeks. By September, second or early third instar larvae descend to the forest litter and overwinter. Larvae resume feeding the following spring, continue developing, pupate and emerge mainly in June – first half of July. This represents a rapid life cycle (Boldaruev, 1955; Rozhkov, 1963). However, under some environmental conditions, a subset of larvae experiences an extended larval stage, prolonging development through the summer months. These larvae descend again in autumn for a second overwintering, thus exhibiting a three-season life cycle (Baranchikov & Kirichenko, 2002). This ontogenetic variability may cause shifts in adult flight timing between different cohorts; however, the long-term data from the south of continental Russian Far East show that the primary driver of variation in flight onset timing is ambient temperature. After correction for the onset of vegetation growth, the timing of flight onset falls within a fairly narrow range. The mathematical result obtained from a substantial sample of seasonal observations demonstrates the applicability of the proposed model to data from sparse populations, where large numbers of zero-value observations are inevitable. This approach resolves the problem of catch variability at low insect population density.

Our study demonstrates that long-term sparse populations of one of the most important phytophages of boreal forests follow the same ecological dynamics as eruptive populations of this or closely related species elsewhere in the range. Understanding the reasons why outbreak potential is constrained in specific regions will provide a foundation for pest population management. Our study also empirically demonstrated, using a large sample, the applicability of binary models to time series containing large numbers of zeros. This enables analysis of population dynamics data for species that spend far more time at low density than in an eruptive phase.

## Acknowledgments

The study was supported by a grant from a state funding program of the Sirius Federal Territory: “Scientific and technological development of the Sirius federal territory” (Agreement No. 24–03 dated 27 September 2024).

## References

Baranchikov YN, Kondakov Y. 1997. Outbreaks of the Siberian moth Dendrolimus superans sibiricus Tschtvrk in Central Siberia, p 10–13. In Gottschalk K, Fosbroke S (ed), Proceedings, Interagency Gypsy Moth Forum. USDA interagency gypsymoth forum, USDA Forest Service, Seattle, Washington, USA.

Barnes W.L., Pagano T.S., Salomonson V.V. Prelaunch characteristics of the Moderate Resolution Imaging Spectroradiometer (MODIS) on EOS-AM1. IEEE Transactions on Geoscience and Remote Sensing, 1998, 36(4), 1088-1100. 9.

Box G. F.P., Jenkins G.M. Time series analysis. Forcasting and control. San Francisco: Holden-Day. 1970. 5553 p.

Delpierre N., Vitasse Y., Chuine I., Guillemot J., Bazot S., Rutishauser T., Rathgeber C. Temperate and boreal forest tree phenology: from organ-scale processes to terrestrial ecosystem models. Annals of Forest Science 2016, v. 73. p.5–25. 10.1007/S13595-015-0477-6.HAL-01306442

EPPO Dendrolimus Sibiricus. EPPO Datasheets on Pests Recommended for Regulation. 2023. Available online: https://gd.eppo.int (accessed on 28 August 2023).

Fischbein D, Corley JC. Population ecology and classical biological control of forest insect pests in a changing world. For Ecol Manag. 2022;520: 120400.

Flø D, Rafoss T, Wendell M, Sundheim L. 2020. The Siberian moth (Dendrolimus sibiricus), a pest risk assessment for Norway. For Ecosyst 7:48. 10.1186/s40663-020-00258-9.

Isaev A.S., Khlebopros R.G., Nedorezov L.V., Kondakov Yu.P., Kiselev V.V., Soukhovolsky V.G. Forest Insects population dynamics. M.: Nauka. 2001. 374 p. (in Russian).

Kharuk VI, Antamoshkina OA. 2017. Impact of silkmoth outbreak on taiga wildfres. Contemp Probl Ecol 10:556–562. 10.1134/S1995425517050055.

Kirichenko, N.; Flament, J.; Baranchikov, Y.; Grégoire, J.C. Native and exotic coniferous species in Europe–possible host plants for the potentially invasive Siberian moth, Dendrolimus sibiricus 1 Tschtv. (Lepidoptera, Lasiocampidae). EPPO Bull. 2008, 38, 259–263.

Kirichenko, N.I.; Ageev, A.A.; Astapenko, S.A.; Golovina, A.N.; Kasparyan, D.R.; Kosheleva, O.V.; Timokhov, A.V.; Tselikh, E.V.; Zakharov, E.V.; Musolin, D.L.; et al. The diversity of parasitoids and their role in the control of the Siberian moth, Dendrolimus sibiricus (Lepidoptera: Lasiocampidae), a major coniferous pest in Northern Asia. Life 2024, 14, 268.

Kondakov Yu. P. Regularities of outbreaks of the Siberian silk moth) // Ecology of Population of animals Siberia. Novosibirsk: Nauka, 1974. P. 206–265 (in Russian)]

Kovalev A., Tarasova O., Soukhovolsky V., Ivanova Yu. D. Is It Possible to Predict a Forest Insect Outbreak? Backtesting Using Remote Sensing Data. Forests 2024, 15(8), 1458. 10.3390/f1508145

Li, W.; Zheng, T.; Yang, Z.; Li, M.; Sun, C.; Yang, X. Classification and detection of insects from field images using deep learning for smart pest management: A systematic review. Ecol. Inform. 2021, 66, 101460.

Økland B, Bjørnstad ON (2006) A resource-depletion model of forest insect outbreaks. Ecology 87:283–290. 10.1890/05-0135

Phenology: An Integrative Environmental Science; Schwartz, M.D., Ed.; Springer: Dordrecht, The Netherlands, 2013; 385 p.

Royama T. Population Dynamics of the Spruce Budworm Choristoneura Fumiferana. Ecological Monographs. 1984;54(4):429–62.

Shimoda, M. & Honda, K.-I. Insect reactions to light and its applications to pest management. Appl. Entomol. Zool. 48, 413–421 (2013).

Shipova A.A., Belousova I. A., Yakimova ME, Kirichenko N.I., Ageev A.A., Cusson M, Lukhtanov V.A., Ershov N.I., Martemyanov V.V. 2026. From Adam’s rib: the Siberian moth’s female W 1 chromosome is derived from its Z counterpart. Molecular Biology and Evolution. Accepted. DOI: 10.1093/molbev/msag055

Soukhovolsky V., Kovalev A., Tarasova O., Martemyanov V. Regulatory characteristics of population density dynamics of forest insects and possible reasons for the observed narrow range of such characteristics. Chaos, Solitons and Fractals 191 (2025) 115949 10.1016/j.chaos.2024.115949

Soukhovolsky V., Kovalev A., Akhanaev Yu, Kurenshchikov D., Ponomarev V., Tarasova O., Caroulle F., Inoue M.N., Martemyanov V. An Autoregulatory Model of Forest Insect Population Dynamics and Forest Stand Damage Dynamics in Different Habitats: An Example of Lymantria dispar L. Forests, 2024. 15(7), 1098. DOI:10.3390/f15071098.

Soukhovolsky V.G., Tarasova O.V., Kovalev A.V., Ivanova Yu.D., Pavlushin S.V., Akhanaev Y.B., Martemyanov V.V Forest insect populations: modeling of critical events as first- and second-order phase transitions. Ecological Modelling. Volume 504, May 2025, 111090 10.1016/j.ecolmodel.2025.111090

Stewart D, Djoumad A, Holden D, Kimoto T, Capron A, Dubatolov VV, Akhanaev Yu B, Yakimova ME, Martemyanov VV, Cusson M. 2023. A Taq-Man assay for the detection and monitoring of potentially invasive lasiocampids, with particular attention to the Siberian Silk Moth, Dendrolimus sibiricus (Lepidoptera: Lasiocampidae). J Insect Science 23:5. 10.1093/jisesa/ieac062.

Teixeira, A.C.; Ribeiro, J.; Morais, R.; Sousa, J.J.; Cunha, A. A systematic review on automatic insect detection using deep learning. Agriculture 2023, 13, 713.

Wai-Ki Ching, Ximin Huang, Michael K. Ng, Tak-Kuen Siu. Markov Chains. Models, Algorithms and Applications. Springer: New York, Heidelberg, Dordrecht, London. 2013. 243 p.

Wei W.W.S. Time series Analysis. Boston: Addison Wesley. 2006. 614 p.

Witzgall, P.; Kirsch, P.; Cork, A. Sex pheromones and their impact on pest management. J. Chem. Ecol. 2010, 36, 80–100.

Wolkovich, E.M.; Cook, B.I.; Davies, T.J. Progress towards an interdisciplinary science of plant phenology: Building predictions across space, time and species diversity. New Phytol. 2014, 201, 1156–1162.

Yurchenko G. I., Turova G. I. Siberian and white-striped silkmoths in the Far East (monitoring manual)). Khabarovsk: Far East For. Res. Inst.), 2007. 98 p. (in Russian)].

Zolotuhin, V. V. 2015. Kokonopryady (Lepidoptera, Lasiocampidae) fauny Rossii i sopredel’nykh territoriy [Lappet Moths (Lepidoptera, Lasiocampidae) of Russia and adjacent territories]. Ulyanovsk: Korporatsiya teckhnologiy prodvizheniya. pp. 384 (in Russian).

